# Functional Tissue Units in the Human Reference Atlas

**DOI:** 10.1101/2023.10.16.562593

**Authors:** Supriya Bidanta, Katy Börner, Bruce W. Herr, Marcell Nagy, Katherine S. Gustilo, Rachel Bajema, Libby Maier, Roland Molontay, Griffin Weber

## Abstract

Functional tissue units (FTUs) form the basic building blocks of organs and are important for understanding and modeling the healthy physiological function of the organ and changes during disease states. In this first comprehensive catalog of FTUs, we document the definition, physical dimensions, vasculature, and cellular composition of 22 anatomically correct, nested functional tissue units (FTUs) in 10 healthy human organs. The catalog includes datasets, illustrations, an interactive online FTU explorer, and a large printable poster. All data and code are freely available. This is part of a larger ongoing international effort to construct a Human Reference Atlas (HRA) of all cells in the human body.

## Introduction

The Human Reference Atlas (HRA) is an international effort of 17 consortia to build a freely available map of the healthy adult human body down to the single-cell level. At the highest level are organ systems, whose individual organs contain various tissues, also called anatomical structures. Tissues, in turn, consist of repeating structures known as functional tissue units (FTUs).

Bernard de Bono et al. define FTUs as “a three-dimensional block of cells centered around a capillary, such that each cell in this block is within diffusion distance from any other cell in the same block”^1^. The description of the cells within an FTU and the anatomical location of the FTU form a so-called primary tissue motif (PTM). An FTU must support both metabolism and communication between cells. The maximum diffusion distance of oxygen and other molecules constrains the size of an FTU. Therefore, tissues and organs in the body consist of many repeating FTUs to perform the overall function of the organ.

We previously generalized this original definition by describing an FTU as “the smallest tissue organization that performs a unique physiologic function and is replicated multiple times in a whole organ”^2^. Here, we extend the definition of FTUs by allowing them to form a nested hierarchy to accomplish various functions.

In this perspective, we present a catalog of the first 22 FTUs for the HRA. The current version of this atlas contains (1) the length and diameter of each FTU; (2) the vasculature pathways that connect FTUs to the heart and amongst each other; (3) the cell types within each FTU; and (4) 2D illustrations of each FTU showing their prototypical cell types, cell shapes, sizes, and spatial arrangement. We map all anatomical structures to their corresponding Uber-anatomy Ontology (UBERON) ID^3^ and all cell types to their Cell Ontology (CL) ID^4^ to enable linkage to other datasets. We reference supporting scholarly paper evidence where applicable. We make the HRA’s catalog of FTUs available as machine-readable downloadable files, through an HRA Interactive FTU Explorer website, and as a large printable poster.

Much of the information we present here about FTUs exists in the literature but siloed by organ or tissue type and with no unifying conceptual or data schema framework. The creation of a catalog of FTUs as an integral part of the HRA, along with standardized descriptions linked to ontologies, makes it possible to compare and contrast these units throughout the human body. This way, we can gain a better comprehension of how FTUs function together, and how they can be integrated into models and simulations.

## HRA FTU Datasets

This initial release of the atlas contains 22 FTUs across 10 organs, see **Table 1**. The 15 top-level FTUs by organ are (1) kidney: nephron; (2) large intestine: crypt of Lieberkuhn of colon; (3) liver: liver lobule; (4) lung: alveolus and bronchus submucosal gland; (5) pancreas: islets of Langerhans, pancreas acini, and intercalated duct; (6) prostate gland: prostate glandular acinus; (7) skin: dermal papilla and skin epidermal ridge; (8) small intestine: intestinal villus; (9) spleen: red pulp and white pulp; and (10) thymus: thymus lobule. The kidney nephron contains 7 additional FTUs: renal corpuscle, inner medullary collecting duct, descending limb of loop of Henle, loop of Henle ascending limb thin segment, thick ascending limb of loop of Henle, outer medullary collecting duct, and cortical collecting duct. **Table 1** includes the UBERON ID, dimensions, and references for each FTU.

**Table 1.** Functional tissue units by organ, with UBERON ID, and dimensions (length where applicable and diameter in millimeters) as listed in the provided references.

| Organ | FTU | UBERON ID | Dimensions | References |
| --- | --- | --- | --- | --- |
| kidney | nephron | UBERON:0001285 | 30-50 | [5] |
| kidney | – renal corpuscle | UBERON:0001229 | 0.15 - 0.25 | [6] |
| kidney | – inner medullary collecting duct | UBERON:0004205 | 12 x 0.05 | [7] |
| kidney | – descending limb of loop of Henle | UBERON:0001289 | 3.2 x 0.026 | [7] |
| kidney | – loop of Henle ascending limb thin segment | UBERON:0004193 | 5 x 0.026 | [7] |
| kidney | – thick ascending limb of loop of Henle | UBERON:0004193 | 5 x 0.026 | [7] |
| kidney | – outer medullary collecting duct | UBERON:0004204 | 5 x 0.040 | [7] |
| kidney | – cortical collecting duct | UBERON:0004203 | 20 - 22 x 0.02 - 0.05 | [7],[8] |
| large intestine | crypt of Lieberkuhn | UBERON:0001983 | 0.05 x 0.7 | [9] |
| liver | liver lobule | UBERON:0004647 | 1 - 2.5 | [10],[11] |
| lung | alveolus of lung | UBERON:0002299 | 0.184 - 0.2 | [12] |
| lung | bronchus submucosal gland | UBERON:8410043 | 1 - 2 | [13] |
| pancreas | islets of Langerhans | UBERON:0000006 | 0.1 | [14] |
| pancreas | pancreas acinus | UBERON:0001263 | 0.010 - 0.024 | [15] |
| pancreas | intercalated duct | UBERON:0014726 | 1.5 - 3.5 | [16] |
| prostate gland | prostate glandular acinus | UBERON:0004179 | 0.5 | [17] |
| skin | dermal papilla | UBERON:0000412 | 0.1 - 1.5 | [18] |
| skin | epidermal ridge of digit | UBERON:0013487 | 0.25–0.93 | [18]–[20] |
| small intestine | villus | UBERON:0001213 | 0.5–1.6 | [22] |
| spleen | red pulp of spleen | UBERON:0001250 | 0.02 - 0.04 | [23] |
| spleen | white pulp of spleen | UBERON:0001959 | 0.5 - 1 | [23] |
| thymus | thymus lobule | UBERON:0002125 | 0.5 - 2 | [20]–[23] |

HRA FTUs are developed in a five-step process: 1) identify FTU shape, dimensions, and cell types from experimental data published in scholarly papers; 2) invite organ experts to comment on these and revise as needed; 3) a professional medical illustrator creates a vector-based drawing of the FTU and saves it as an SVG file with all metadata; 4) organ experts review the drawings and any existing disclaimers and suggest changes as needed; 5) number of cells per cell type are recorded and the FTU file is published with all metadata together with a crosswalk file as part of an HRA release. Subsequently, we detail key data types and formats.

### Geometric Properties

**Table 1** lists the 10 organs and their 22 FTUs together with the UBERON ID, spatial dimensions, and reference paper(s) in which the dimensions were published. For circular-shaped FTUs, such as the *alveolus of the lung*, a single dimension value (or range) is listed representing the typical diameter of the FTU. For cylindrical FTUs, such as the *inner medullary collecting duct* in the kidney, both length and diameter are provided. The diameter of the *intercalated duct* is the largest at the head of the pancreas and smallest in the tail.

### Vasculature

The HRA-VCCF dataset contains a list of all the blood vessels in the HRA, along with their branching structure, cell types, biomarkers, and other information^5,6^. **Table 2** lists the vessels in the HRA-VCCF that directly supply or drain each FTU. Note that the epidermal ridge does not contain blood vessels, but rather obtains oxygen via diffusion from the underlying dermal papilla. **Supplemental Table S1** lists the full vasculature pathways from the heart to each FTU and back to the heart. Where possible, vessels are mapped to their corresponding UBERON ID or Foundational Model of Anatomy (FMA) ontology IDs ^7^. About 63% of the vessels exist in one or both of these ontologies.

**Table 2.** Blood vessels that directly supply or drain each FTU.

| Organ | FTU | Vessels |
| --- | --- | --- |
| kidney | nephron | ascending vasa recta of kidney; descending vasa recta of kidney; glomerular capillary; peritubular capillary; renal afferent arteriole; renal efferent arteriole; vasa recta of kidney |
| kidney | – cortical collecting duct | peritubular capillary |
| kidney | – descending limb of loop of Henle | ascending vasa recta of kidney |
| kidney | – inner medullary collecting duct | ascending vasa recta of kidney; descending vasa recta of kidney |
| kidney | – loop of Henle ascending limb thin segment | vasa recta of kidney |
| kidney | – outer medullary collecting duct | ascending vasa recta of kidney; descending vasa recta of kidney |
| kidney | – renal corpuscle | glomerular capillary; renal afferent arteriole; renal efferent arteriole |
| kidney | – thick ascending limb of loop of Henle | peritubular capillary |
| large intestine | crypt of Lieberkuhn | branch of mucous plexus of colon |
| liver | liver lobule | central vein of liver; hepatic arteriole; hepatic portal venule; hepatic sinusoid |
| lung | alveolus of lung | alveolar capillary |
| lung | bronchus submucosal gland | bronchial capillary |
| pancreas | intercalated duct | periductal capillary of intercalated duct of pancreas |
| pancreas | islet of Langerhans | pancreatic islet capillary |
| pancreas | pancreatic acinus | intralobular arteriole of pancreas; lobular capillary of pancreas |
| prostate gland | prostate glandular acinus | prostatic capillary |
| skin | dermal papilla | branch of subpapillary plexus |
| skin | epidermal ridge of digit | (indirectly via branch of subpapillary plexus in the dermal papilla) |
| small intestine | intestinal villus | branch of mucous plexus of small intestine |
| spleen | red pulp of spleen | post-sheath open capillary of spleen; red pulp venule of spleen; splenic venous sinusoid |
| spleen | white pulp of spleen | branch of superficial white pulp capillary of spleen; secondary follicle arteriole of spleen; spleen central arteriole |
| thymus | thymus lobule | thymic capillary |

### Cell Types

A list of the different types of cells in each FTU is available at https://humanatlas.io/assets/table-data/ftu-cell-count-5th-release.csv.

**Table 3** shows a subset of this list for FTUs within the kidney nephron. Each cell type is associated with its corresponding Cell Ontology (CL) ID. Ongoing research using methods such as single-cell RNA sequencing is identifying biomarkers expressed in these cell types and determining the relative distribution of the cell types in the FTUs. Examples for the liver^8^ and lung^9^ are available at https://www.ebi.ac.uk/gxa/sc/experiments/E-MTAB-10553/results/anatomogram and https://www.ebi.ac.uk/gxa/sc/experiments/E-GEOD-130148/results/anatomogram.

**Table 3.** Cell types with CL IDs in kidney nephron FTUs.

| FTU | Cell Type | CL ID |
| --- | --- | --- |
| renal corpuscle | parietal epithelial cell | CL:1000452 |
| renal corpuscle | glomerular visceral epithelial cell | CL:0000653 |
| renal corpuscle | glomerular capillary endothelial cell | CL:1001005 |
| renal corpuscle | glomerular mesangial cell | CL:1000742 |
| renal corpuscle | epithelial cell of proximal tubule | CL:0002306 |
| renal corpuscle | macula densa epithelial cell | CL:1000850 |
| renal corpuscle | kidney afferent arteriole endothelial cell | CL:1001096 |
| renal corpuscle | kidney efferent arteriole endothelial cell | CL:1001099 |
| cortical collecting duct | kidney cortex collecting duct principal cell | CL:1000714 |
| cortical collecting duct | kidney cortex collecting duct intercalated cell type A | CL:4030020 |
| cortical collecting duct | kidney cortex collecting duct intercalated cell type B | CL:4030021 |
| cortical collecting duct | peritubular capillary endothelial cell | CL:1001033 |
| inner medullary collecting duct | kidney inner medulla collecting duct principal cell | CL:1000718 |
| inner medullary collecting duct | peritubular capillary endothelial cell | CL:1001033 |
| outer medullary collecting duct | kidney outer medulla collecting duct intercalated cell type A | CL:1000717 |
| outer medullary collecting duct | peritubular capillary endothelial cell | CL:1001033 |
| outer medullary collecting duct | kidney outer medulla collecting duct principal cell | CL:1000716 |
| thick ascending loop of Henle | kidney loop of Henle thick ascending limb epithelial cell | CL:1001106 |
| thick ascending loop of Henle | peritubular capillary endothelial cell | CL:1001033 |
| descending thin loop of Henle | kidney loop of Henle thin descending limb epithelial cell | CL:1001111 |
| descending thin loop of Henle | vasa recta descending limb | CL:1001285 |
| ascending thin loop of Henle | kidney loop of Henle thin ascending limb epithelial cell | CL:1001107 |
| ascending thin loop of Henle | vasa recta ascending limb | CL:1001131 |

### Spatial Arrangement of Cells

The spatial arrangement of cells and their types in each FTU is recorded in 2D illustrations, see https://humanatlas.io/2d-ftu-illustrations.

Medical illustrators create these illustrations by compiling information from papers, histology, and microscopical images of the FTU guided by Standard Operating Procedures (SOP)^3^ and the *Style Guide for Human Reference Atlas 2D Functional Tissue Unit (FTU) Illustrations*^10^. The information is validated by Subject Matter Experts (SMEs) with extensive expertise in human anatomy and single-cell studies. The illustrations are saved in SVG format, and converted to JSON files using a code that helps to map the drawings to the metadata; and a Crosswalk table is compiled that associates 2D anatomical structures and cell types in the FTUs with their proper terms in the HRA. Illustrations of all 22 FTUs are shown in **Fig. 1** at four levels of magnification, see scale bars.

**Figure 1.**
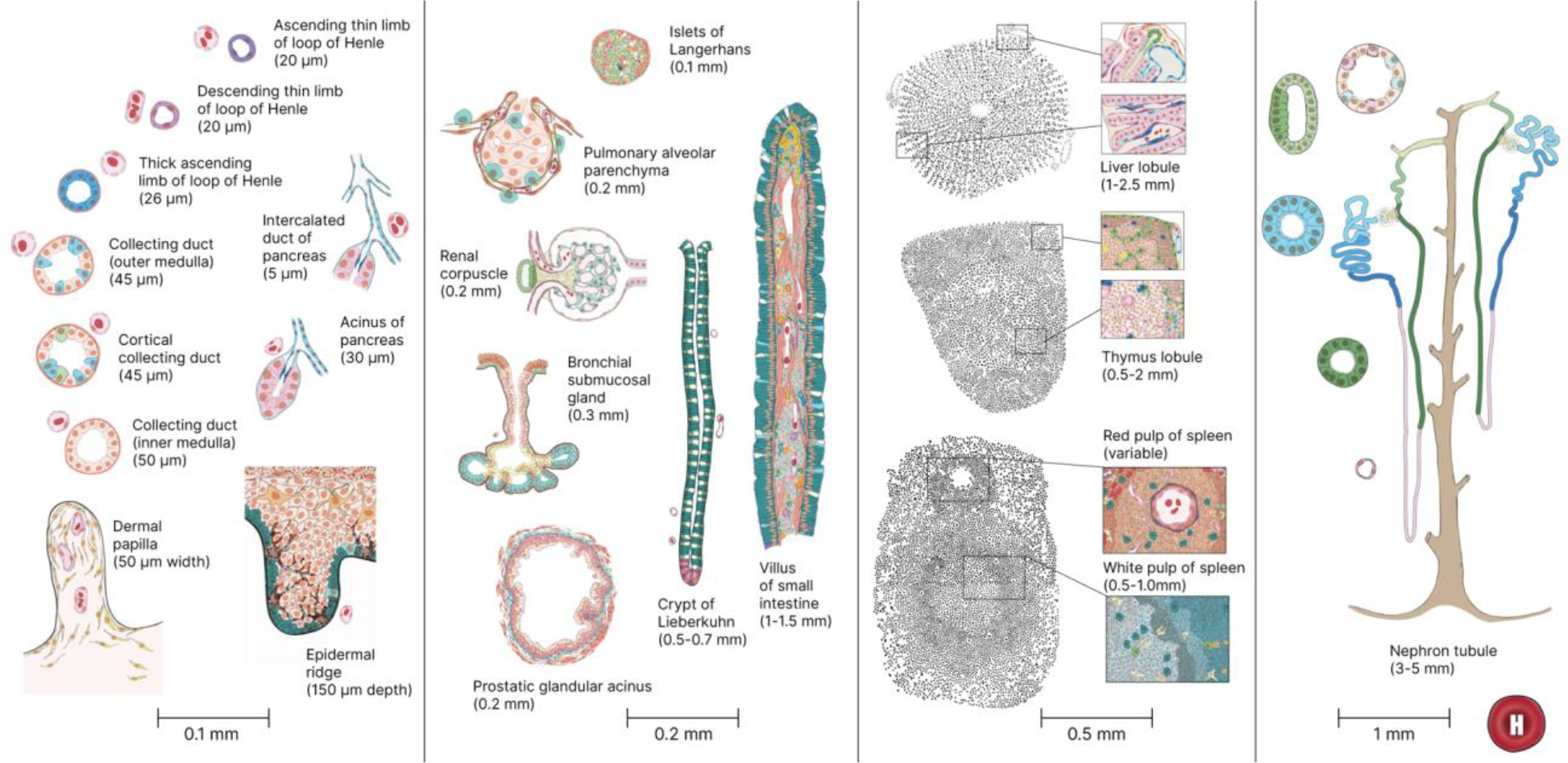
2D illustrations of all 22 FTU with name and size annotations.

## HRA FTU Visualizations

In addition to accessing the HRA FTUs through machine-readable data files, users can also explore the data online using the Interactive FTU Explorer or as a printable poster.

### Interactive FTU Explorer

Prior work by the European Bioinformatics Institute (EBI) on interactive anatomograms^11,12^ and by the Kidney Precision Medicine Project (KPMP) on the Kidney Tissue Atlas Explorer^13^ inspired the design of the lung and kidney FTU illustrations (creating de-facto standards across these three sites) and the design and implementation of the HRA Interactive FTU Explorer at https://hubmapconsortium.github.io/hra-ui/apps/ftu-ui/.

As illustrated in **Fig. 2**, the website lets users select any of the 22 FTUs on the left, which brings up an interactive rendering of the FTU illustration in the middle. Cell types and their biomarkers derived from experimental data are tabulated in the top right; with circle size indicating the percentage of cells in the FTU and circle color representing the mean expression value over all cells of this type in the FTU—averaged over all datasets loaded for this FTU. Users can choose to explore gene, protein, and lipid biomarkers for different cell types using three tabs; in some cases, experimental data might not (yet) exist. In the lower right of the FTU Explorer, users can click on a specific source dataset and go to the portal that serves this dataset. In the next iteration of the FTU Explorer, users will be able to select specific datasets (e.g., datasets from different demographics) to understand changes in cell type by biomarker expression values as we age (young vs. old) or by sex (male vs. female).

**Figure 2:**
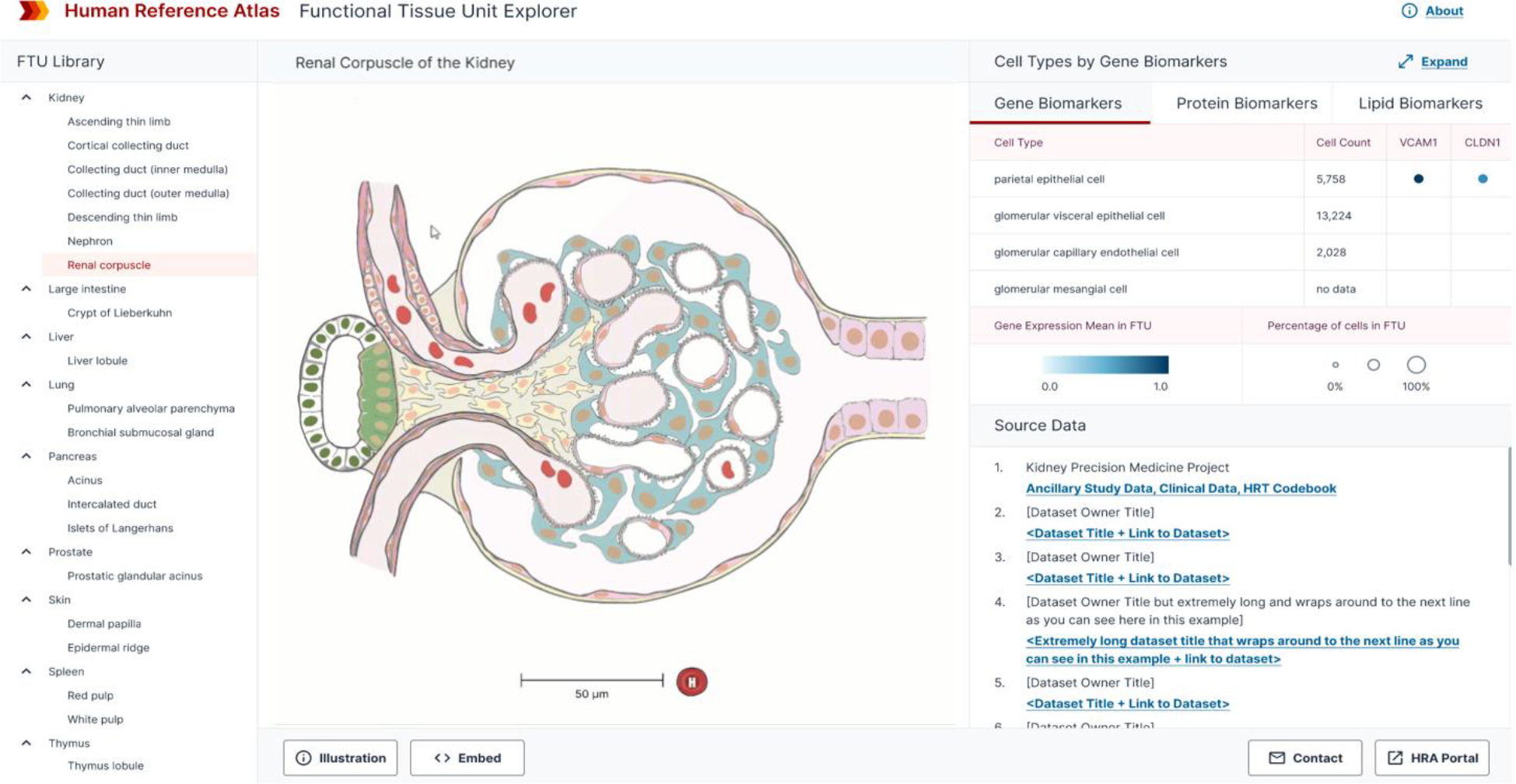
HRA Interactive Functional Tissue Unit Explorer showing the 2D illustration of the renal corpuscle of the kidney along with associated experimental data.

### Printable Poster

To place the FTUs within the larger context of the HRA, we created a printable poster visualizing all 1,607 anatomical structures and 1,943 cell types available in the 5th release of the HRA. The visualization is composed of two radial tree graphs: (1) The first graph contains the nested “partonomy” of the anatomical structures and cell types in the HRA. The human body serves as the root node in the center, the largest anatomical structures (organs) are placed further out, and smaller sub-structures and tissue types of branch outwards from the organs’ leaf nodes denoting cell types. (2) The second graph contains all the blood vessels in the HRA, with the chambers of the heart in the center, and increasing smaller vessels more distal to the heart again branching outwards from the center. Nodes in the two radial tree graphs meet at points where the HRA indicates a vessel supplies or drains the corresponding anatomical structure. Nodes that represent FTUs are highlighted in green. The left side shows the nested partonomy and blood vessels in females and the right side shows them for males with a “butterfly-like” appearance that invites closer examination, discussion, and self-portrait photos, see **Fig. 3**. The 6-foot diameter poster is available in ready-to-print formats via https://github.com/cns-iu/hra-vccf-ftu-supporting-information.

**Figure 3.**
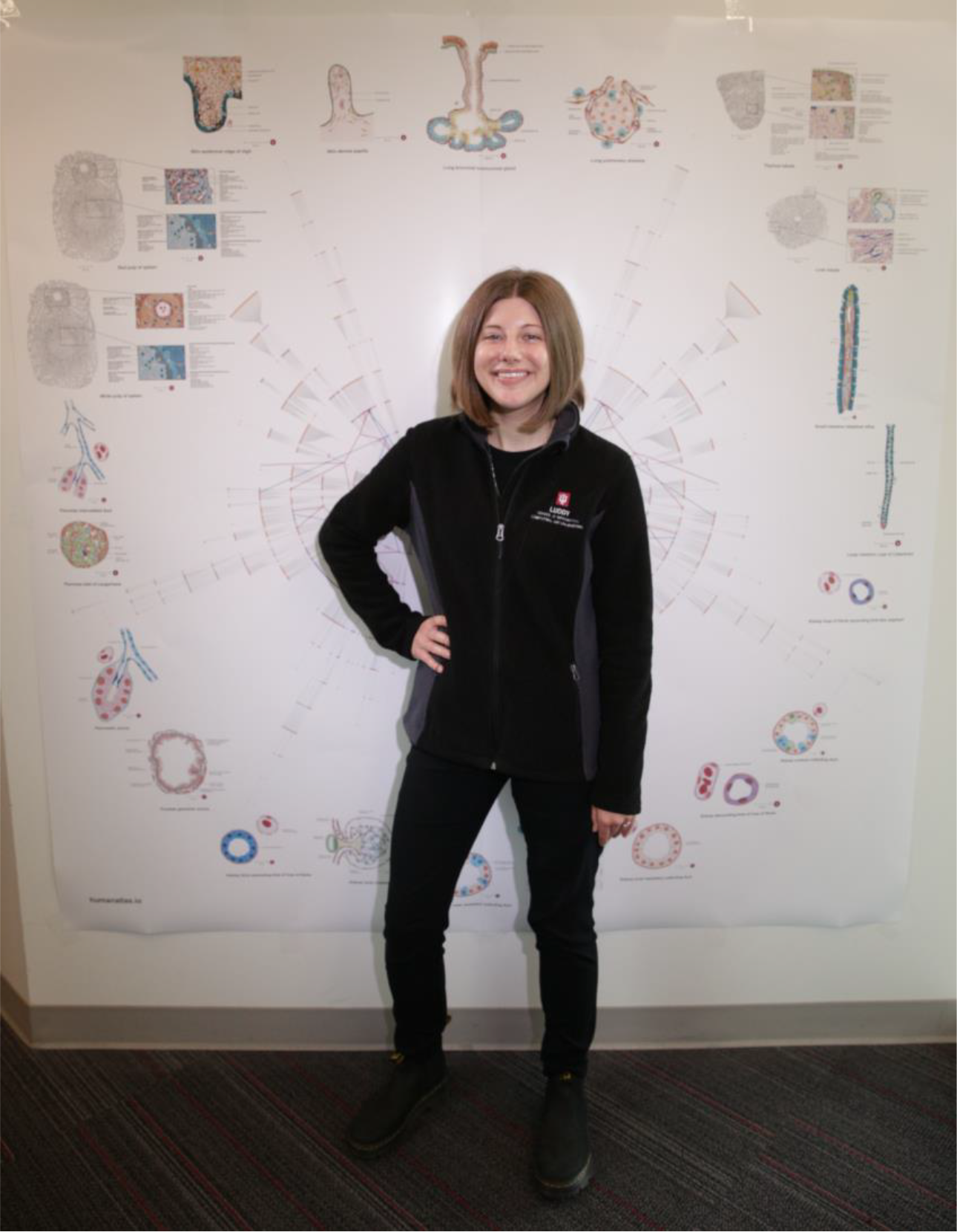
Poster of the HRA illustrating radial tree graphs of (1) the nested partonomy of organ anatomical structures and cell types of the human body with an overlay of (2) the branching structure of the blood vasculature extending from the heart (center of the figure) to FTUs (outer edge). Illustrations of all 22 FTUs are placed outside of the radial tree visualization for easy reference. One of the authors is posing for a photo.

Note that there are 54 nodes for the 22 FTUs. This is because some FTUs overlap multiple anatomical structures and therefore appear in more than one location of the partonomy. We plan to update this poster with future releases of the HRA to reflect ongoing development of the partonomy and vasculature graphs and new FTUs added to the catalog.

## Discussion

FTUs are the basic building blocks of organs. Their distinctive size (relative to diffusion distances) and physical arrangement of different cell types are key to enabling their corresponding physiologic function. In this perspective, we described datasets and visualizations of 22 FTUs in 10 organs interconnected by vasculature as published in the 5th release of the HRA. To our knowledge, this is the first time that comprehensive data specifically about FTUs across the human body have been systematically cataloged. At the macro-anatomical scale, the vascular pathways connecting FTUs to the heart and each other, along with our HRA poster, show the position of FTUs in the body. At the microscopic scale, the illustrations and cell type table show the internal structure of the FTUs.

Within the HRA effort, we are using this catalog of FTUs for several use cases. For example, we ran Kaggle competitions to develop scalable and generalizable segmentation code to identify FTUs in images^2,14^. We use the segmentation code to automatically count the number of FTUs per unit area and to compute general biomarker expression values for genes, proteins, lipids, and metabolites. Ongoing research compares hierarchical cell neighborhoods^15^ computed from tissue data with the 22 FTUs in the catalog presented here.

More broadly, we envision the FTU catalog and the Interactive FTU Explorer as a framework for researchers to study how the information presented here (FTU dimensions, cell types, etc.) varies across donor demographics (e.g., age, sex, race) and in different disease states. Understanding similarities and differences between FTUs can help predict adverse events of medications or suggest new drug targets. The FTU Explorer can be used as an educational tool, especially for comparing the physiology and vasculature of different organs and tissues. Our downloadable poster is intended for general public outreach.

A limitation of this work is that the HRA is still in development, with updates planned every six months for the coming three years. As a result, this is not yet a complete map of all FTUs in the body, but rather a starting point demonstrating how various types of data about FTUs, including physical dimensions, vasculature, cell types, and spatial orientation can be interlinked across scales, in support of creating a human reference atlas.

## Data availability

All data is freely available at https://github.com/cns-iu/hra-vccf-ftu-supporting-information and via the HRA Portal at https://humanatlas.io.

## Code availability

All code is freely available at https://github.com/cns-iu/hra-vccf-ftu-supporting-information. The FTU Explorer code is hosted at https://github.com/hubmapconsortium/hra-ui. Standard operating procedures are available via the HRA Portal, https://humanatlas.io. HRA learning modules and self-quizzes can be accessed via the Visible Human MOOC at https://expand.iu.edu/browse/sice/cns/courses/hubmap-visible-human-mooc.

## Acknowledgments

We thank organ experts Sanjay Jain, Matthias Kretzler, M. Todd Valerius (Kidney), John Hickey, Yiing Lin (Intestine), Gloria Pryhuber (Lung), Martha Campbell-Thompson (Pancreas), Douglas Strand (Prostate), Fiona Ginty, Arivarasan Karunamurthy (Skin), Andrea J. Radtke, Maigan Brusko (Spleen), Maigan Brusko, Andrea J. Radtke (Thymus), Anna Maria Masci, Tim Kendall, Ayako Suzuki (Liver) who supported FTU designs and revisions. Discussions with Michael Rose, Todd M. Valerius, and Matthias Kretzler from the KPMP team working on the Kidney Tissue Atlas Explorer and Silvie Fexova and Irene Papatheodorou from EBI working on the Anatomogram Explorer informed the design of the Interactive FTU Explorer user interface. The FTU Explorer was implemented by Daniel Bolin, Edward Lu, Raj Chavan, Rohith Reddy, Sai Kiran Jella, Himanshu Joshi, Sanket Darwante, Jagrut Chaudhari, Sumedh Salvi, Pranathi Neerudu, Anita Narayanan, Aditya Rudrawar, Tanu Kansal, Hanuhitha Dondapati, and Akshay Murthy. Heidi Schlehlein designed the butterfly visualization poster using files provided by the authors of this paper. Ellen Quardokus helped in validating the cell types and provided guidance for finalizing the FTU data.

This research has been funded by the NIH Common Fund through the Office of Strategic Coordination/Office of the NIH Director under awards OT2OD033756 and OT2OD026671, by the Cellular Senescence Network (SenNet) Consortium through the Consortium Organization and Data Coordinating Center (CODCC) under award number U24CA268108, by the Kidney Precision Medicine Project grant U2CDK114886, by the NIDDK under awards U24DK135157 and U01DK133090 and by The Multiscale Human CIFAR project. The funders had no role in study design, data collection and analysis, decision to publish, or preparation of the manuscript.

## Author contributions

SB compiled FTU data; KB lead this work as part of the HRA effort; MN and RM implemented the butterfly visualization; KSG and GW compiled the vasculature data; RB designed the FTU illustrations with expert input by organ experts; LM, BH, and KB developed the design specification for the FTU Explorer; BH led FTU Explorer development; SB, MN, KB, and GW wrote the manuscript. All authors edited the manuscript.

## Supplemental table

**Table S1**. Vascular pathways from the heart to each FTU and back to the heart https://github.com/cns-iu/hra-vccf-ftu-supporting-information/tree/main/data

## References

1. De Bono, B., Grenon, P., Baldock, R. & Hunter, P. Functional tissue units and theirprimary tissue motifs in multi-scale physiology. J. Biomed. Semant. 4, 22 (2013).

2. Jain, Y. et al. Segmenting functional tissue units across human organs using community-driven development of generalizable machine learning algorithms. Nat. Commun. 14, 4656(2023).

3. Börner, K. et al. Anatomical structures, cell types and biomarkers of the HumanReference Atlas. Nat. Cell Biol. 23, 1117–1128 (2021).

4. Osumi-Sutherland, D. et al. Cell type ontologies of the Human Cell Atlas. Nat. Cell Biol.23, 1129–1135 (2021).

5. Nephron | Definition, Function, Structure, Diagram, & Facts | Britannica.https://www.britannica.com/science/nephron (2023).

6. Physiology Image: Renal corpuscle. - PhysiologyWeb.https://www.physiologyweb.com/figures/physiology_image_TvKcIccyvFuiIvXAWDRXu5GuzkBK8RHt_renal_corpuscle.html.

7. Layton, A. T. & Layton, H. E. A computational model of epithelial solute and watertransport along a human nephron. PLOS Comput. Biol. 15, e1006108 (2019).

8. Himmerkus, N. et al. Viewing Cortical Collecting Duct Function Through Phenotype-guided Single-Tubule Proteomics. Function 1, zqaa007 (2020).

9. Wang, Y. et al. Capture and 3D culture of colonic crypts and colonoids in a microarrayplatform. Lab. Chip 13, 4625–4634 (2013).

10. ajeyaseelan. 2i. The Liver. Collection at Bartleby.com https://www.bartleby.com/lit-hub/anatomy-of-the-human-body/2i-the-liver (2022).

11. Gray, H., Gray, H. & Lewis, W. H. Anatomy of the human body. (Lea & Febiger, 1918).doi:10.5962/bhl.title.20311.

12. Ochs, M. et al. The number of alveoli in the human lung. Am. J. Respir. Crit. Care Med.169, 120–124 (2004).

13. Ostedgaard, L. S. et al. Lack of airway submucosal glands impairs respiratory hostdefenses. eLife 9, e59653 (2020).

14. Da Silva Xavier, G. The Cells of the Islets of Langerhans. J. Clin. Med. 7, 54 (2018).

15. Morgan, R. G., Schaeffer, B. K. & Longnecker, D. S. Size and number of nuclei differ innormal and neoplastic acinar cells from rat pancreas. Pancreas 1, 37–43 (1986).

16. R, R. & Weerakkody, Y. Pancreatic duct diameter. in Radiopaedia.org (Radiopaedia.org,2013). doi:10.53347/rID-24634.

17. Fullwood, N. J., Lawlor, A. J., Martin-Hirsch, P. L., Matanhelia, S. S. & Martin, F. L. Ananalysis of benign human prostate offers insights into the mechanism of apocrine secretion andthe origin of prostasomes. Sci. Rep. 9, 4582 (2019).

18. Lambert, P. H. & Laurent, P. E. Intradermal vaccine delivery: Will new delivery systemstransform vaccine administration? Vaccine 26, 3197–3208 (2008).

19. Epidermis (Outer Layer of Skin): Layers, Function, Structure. Cleveland Clinichttps://my.clevelandclinic.org/health/body/21901-epidermis.

20. Lundström, R., Dahlqvist, H., Hagberg, M. & Nilsson, T. Vibrotactile and thermal perception and its relation to finger skin thickness. Clin. Neurophysiol. Pract. 3, 33–39 (2018).

21. Fruhstorfer, H., Abel, U., Garthe, C.-D. & Knüttel, A. Thickness of the stratum corneum of the volar fingertips. Clin. Anat. 13, 429–433 (2000).

22. Intestinal villus. Wikipedia (2023).

23. Elmore, S. A. Enhanced histopathology of the spleen. Toxicol. Pathol. 34, 648–655(2006).

24. Thymus histology. Kenhub https://www.kenhub.com/en/library/anatomy/histology-of-the-thymus.

25. Lobules of thymus - e-Anatomy - IMAIOS. https://www.imaios.com/en/e-anatomy/anatomical-structure/lobules-of-thymus-1553795620.

26. Palumbo, C. Embryology and Anatomy of the Thymus Gland. in Thymus Gland Pathology (eds. Lavini, C., Moran, C. A., Morandi, U. & Schoenhuber, R.) 13–18 (Springer Milan, 2008). doi:10.1007/978-88-470-0828-1_2.

27. Boppana, A. et al. Anatomical structures, cell types, and biomarkers of the healthy human blood vasculature. Sci. Data 10, 452 (2023).

28. Griffin Weber, Yingnan Ju, & Katy Börner. Considerations for Using the Vasculature as aCoordinate System to Map All the Cells in the Human Body. Front. Cardiovasc. Med. 7, (2020).

29. Foundational Model of Anatomy | Structural Informatics Group.http://si.washington.edu/projects/fma.

30. Wang, Z.-Y. et al. Single-cell and bulk transcriptomics of the liver reveals potential targets of NASH with fibrosis. Sci. Rep. 11, 19396 (2021).

31. Vieira Braga, F. A. et al. A cellular census of human lungs identifies novel cell states inhealth and in asthma. Nat. Med. 25, 1153–1163 (2019).

32. Bajema, R. Style Guide: Human Reference Atlas 2D Functional Tissue Unit (FTU)Illustrations. (2022) doi:10.5281/ZENODO.6703376.

33. Experiment < Single Cell Expression Atlas < EMBL-EBI.https://www.ebi.ac.uk/gxa/sc/experiments/E-MTAB-10553/results/anatomogram.

34. Moreno, P. et al. Expression Atlas update: gene and protein expression in multiple species. Nucleic Acids Res. 50, D129–D140 (2022).

35. Explorer. https://atlas.kpmp.org/explorer/.

36. Jain, Y. et al. Segmentation of human functional tissue units in support of a Human Reference Atlas. Commun. Biol. 6, 717 (2023).

37. Hickey, J. W. et al. Organization of the human intestine at single-cell resolution. Nature619, 572–584 (2023).

38. Ginzberg, M. B., Kafri, R. & Kirschner, M. Cell biology. On being the right (cell) size.Science 348, 1245075 (2015).

